# Improved protein model quality assessments by changing the target function

**DOI:** 10.1101/270678

**Authors:** Karolis Uziela, David Menéndez Hurtado, Nanjiang Shu, Björn Wallner, Arne Elofsson

## Abstract

Protein modeling quality is an important part of protein structure prediction. We have for more than a decade developed a set of methods for this problem. We have used various types of description of the protein and different machine learning methodologies. However, common to all these methods has been the target function used for training. The target function in ProQ describes the local quality of a residue in a protein model. In all versions of ProQ the target function has been the S-score. However, other quality estimation functions also exist, which can be divided into superposition- and contact-based methods. The superposition-based methods, such as S-score, are based on a rigid body superposition of a protein model and the native structure, while the contact-based methods compare the local environment of each residue. Here, we examine the effects of retraining our latest predictor, ProQ3D, using identical inputs but different target functions. We find that the c ntact-based methods are easier to predict and that predictors trained on these measures provide some advantages when it comes to identifying the best model. One possible reason for this is that contact based methods are better at estimating the quality of multi-domain targets. However, training on the S-score gives the best correlation with the GDT_TS score, which is commonly used in CASP to score the global model quality. To take the advantage of both of these features we provide an updated version of ProQ3D that predicts local and global model quality estimates based on different quality estimates.

## Introduction

Protein quality estimations has a long history. In the 1990s knowledge-based energy functions using the local environment or residue-residue contacts were developed^1, 2^. These methods were developed to detect the native structure, or to detect the correct fold by threading a sequence through all known protein folds^3^. Later models based on atom-atom contacts were also developed^4^.

Although these methods were quite good at identifying errors in protein structures, they only had limited success when used for model quality estimation. One reason for this is that these energy functions are developed to distinguish between the native structure and an unfolded protein. However, incorrect models share many features of the native structure—making it difficult to distinguish them from the native structure using these energy functions.

To solve this problem model quality estimation methods were developed. These methods are often developed using a machine learning approach trained not only to detect the native structure but to actually predict the quality of a model. One of the first methods using this methodolgy was ProQ^5^.

For more than a decade we have developed protein model quality assessment methods using various input features^6, 5, 7^. The ProQ methods have consistently been among the best individual protein model quality assessment methods in CASP evaluations^8, 9, 10^. During the last few years we have been able to provide a small, but significant, improvement to the methods by (i) including additional descriptions of a protein model ^11^ and (ii) applying deep learning techniques ^12^. Although the overall prediction accuracy has improved substantially the ability to identify the best model has not improved significantly for some time.

When developing a model quality assessment method it is necessary to define the target function to be used. All ProQ as well as many other methods have been trained using the same target function, the S-score ^13, 14^. The S-score is calculated after a superposition of the model and the native structure and then describes the displacement of each residue. A global S-score can then be calculated as the average S-score over a model. The global score is similar in nature to other widely used measures, including the main target function in CASP, GDT_TS ^15, 8^, MaxSub ^16^ and TM-score ^17^. The main difference between these methods is how the sum of the scores is normalised by model and protein target length.

During the last few years contact-based approaches to model quality assessments have been introduced; ModlDDT^18^, CAD-score^19^. These functions have also been used for consensus based MQAPs^20, 21, 22^. All these methods assess the conservation of local interactions between a model and the native structure, i.e. they are not dependent on a global superposition, making the contact-based methods insensitive to variability in relative domain arrangements. This in turn might make them more suitable as target functions for single-model MQAPs that are using the local environment of individual residues as their main descriptor.

Here, we first set out to examine the relationship between the different target functions for model quality assessment. Then, we retrained ProQ3D using the different measures as target functions and evaluated their performance. We show that, as expected, the version of ProQ3D that is trained on one target function also performs best when that target function is used for evaluation. However, when examining the ability to identify the best model out of a set of candidates, it is clear that training ProQ3D on contact-based functions is advantageous.

## Methods

Below, we describe briefly the different model quality estimation methods used in this paper, for complete descriptions see the original papers.

### Superposition-based quality estimators

Superposition-based estimators are using a structural alignment of a model to the native structure. In contrast to traditional measures of the Root Mean Square Deviations (RMSD) the goal of these methods are to only superpose the most similar regions of the two structures. The different superposition-based methods differ on the exact algorithm to identify the most similar region and on the exact target function.

We have used three different superposition-based quality estimators for training and testing: S-score ^14, 13^, TM-score ^23^ and GDT_TS ^24^. The GDT_TS measure was only used for testing, because ProQ3D is trained on the local (residue) level while GDT_TS is defined only on the global (protein) level.

The quality estimator S-score used in ProQ^7, 11, 12, 25^ is defined as:

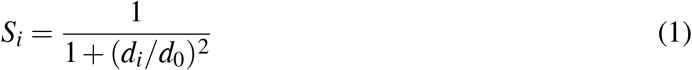

where *d_i_* is the distance between residue i in the model and in the native structure after superposition. Here, the distance threshold *d*_0_ is set to 3Å. The global S-score for the whole model is the sum of all the local scores divided by the protein length.

TM-score is using the same equation (1) as in S-score but the *d*_0_ parameter in TM-score depends on the protein length (*L_N_*) and is defined as:

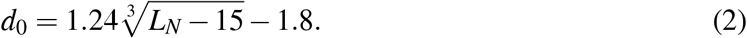

By default, TM-score does not provide the local (per-residue) quality estimates. Therefore, we have run TM-score with the −○ parameter that outputs the structural superposition between the model and the native structure and then calculated the local TM-scores using the formula (1) replacing *d*_0_ with the expression in (2).

GDT_TS was calculated the same way as in the CASP experiments using the LGA package^15^. The program was run with the following parameters:

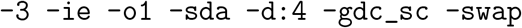

### Contact-based quality estimators

In contrast to superposition-based quality estimators the contact-based predictors do not use a superposition of a model and the native structure. Instead these methods use the local environment, e.g. described by contacts or residue surface areas, for each residue as a descriptor of the quality.

Two contact-based quality estimators were used: lDDT ^18^ and CAD-score ^19^. lDDT compares atom-atom distances between the model and the native structure, while CAD-score compares residue contact surface areas. Both lDDT and CAD-score were run with default parameters, but ignoring residue name consistency checks (`-x` parameter in lDDT and `-b` parameter in CAD-score). For CAD-score we have used the “all atom-sidechain” (A-S) contact score. The reason is that this score behaves better for models with missing residues.

Both CAD and lDDT utilizes all atoms in a protein model while the superposition-based ones only use the *Cα*-coordinates. To examine if the improved performance in first rank loss is due to the fact that CAD and lDDT uses full atom representation we also trained a version of ProQ3D using the version of lDDT that only used the *Cα* atoms (lDDT-CA^18^) as a target function.

### Training deep learning method

We trained the deep learning methods in exactly the same way as in ProQ3D^12^ using different local quality estimators, S-score, a local version of TM-score, lDDT or CAD-score as target functions. For details of the target function see the Methods section. Briefly, ProQ3D uses a multilayer perceptron with two dense hidden layers (600 and 200 neurons). The optimisation function used was Adadelta with a 10^−11^ penalty for the *L*^2^ regularisation on the weights. Training was done using Keras^26^ library with Theano^27^ backend in Python.

Below, we will refer to the different predictors as: ProQ3D-S, ProQ3D-TM, ProQ3D-CAD and ProQ3D-lDDT, when trained on S-score, TM-score, CAD-score and lDDT, respectively, as target functions. Proq3D-S is obviously identical to the original ProQ3D but we use the name ProQ3D-S for clarity.

### Datasets

We have used exactly the same training and test data sets as in ProQ3D ^12^. The training set consisted of CASP9 and CASP10 models and the two test sets consisted of CASP11 and CAMEO^28^ (2014-06-06–2015-05-30) models. Cancelled targets and targets shorter than 50 residues were filtered out. For simplicity we choose to show CAMEO data in the main text and CASP11 data in the supplementary material, as the results are very similar.

To account for sub-optimal side-chain orientations, we have repacked the side-chains of all models as described in the development of ProQ3^11^. Models with repacked side-chains were used for all four predictors: ProQ3D-S, ProQ3D-TM, ProQ3D-CAD and ProQ3D-lDDT.

### Evaluation

We used Pearson correlation for evaluating the agreement between the predicted values and the target values. The correlations are evaluated on local (residue) level and global (model) level. The global correlations are calculated in two ways: the correlation for the whole data set and the average correlation calculated for each target separately. We evaluated both direct correlations, i.e. when the method is trained and tested on the same quality measures, and indirect correlations when the method is trained on one quality measure but tested on another.

Model selection was assessed by the *first ranked loss* ^9^, defined as the average difference between the model with the best quality and the selected model according to a given quality estimator.

Contact order is the average sequence separation between residues in contact in the structure. It was calculated by Perl script available at http://depts.washington.edu/bakerpg/contact_order/^29^. The script was run with `-r` parameter that returns the relative contact order and `-c 6` parameter that sets 6 Å cutoff for contacting residues (default settings).

## Results and Discussion

Here, we study the effect of the choice of a target function for protein model quality estimation. We use five different protein quality evaluation estimates, three based on superposition (GDT_TS, TM-score and S-score) and two superposition independent contact-based methods (CAD-score and lDDT). All methods except GDT_TS provide both local and global quality estimates.

### Correlation between quality estimates

First, we examine the correlation between the four measures that provide local quality estimate (Figure 1). All quality measures correlate with each other. However, it is also clear that the measures using similar principles, i.e. superposition (S-score and TM-score) or contacts (CAD-score and lDDT) are more similar to each other (*cc* = 0.88 and *cc* = 0.79). The correlations between the two groups are lower (*cc* = 0.52 to 0.76). In particular CAD-score only shows a weak correlation with the superposition-based methods.

**Figure 1:**
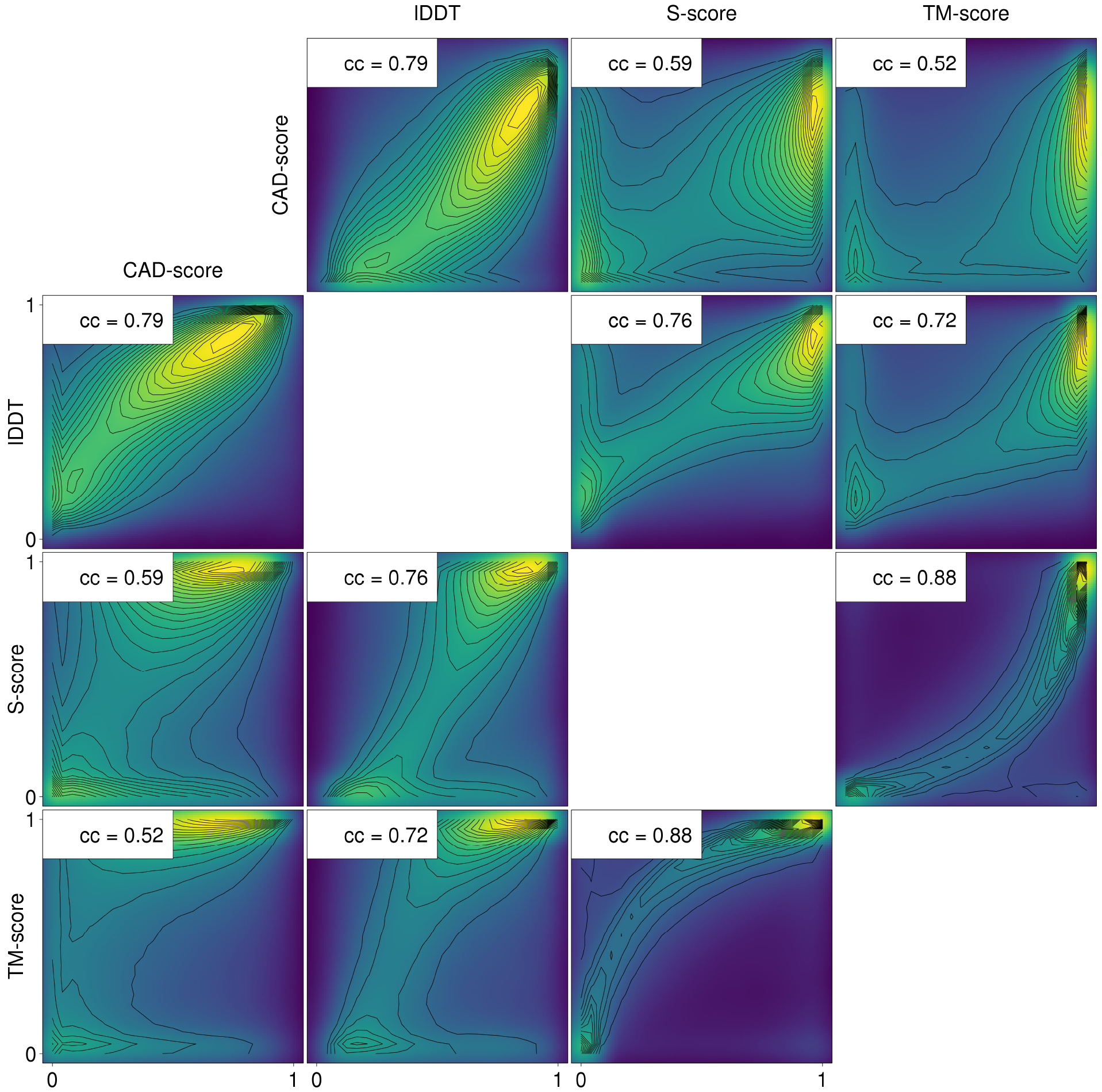
Scatter plot comparing different quality estimators on the *local* level using 4,939,249 residues (points) from models submitted to CAMEO. The intensity is plotted using the Viridis color scale, i.e. the most dense regions are yellow. Contour lines based on the point density are marked. The correlation coefficient (cc) between the different quality estimators i s given in the insets.

It can also be noted that the quality estimated from both TM-score and S-score is almost bimodal with a clear majority of the residues evaluated as good (quality close to one) or bad (quality close to zero). This is because both these measures are using Eq. 1 to calculate the score, effectively assigning a score close to one to most superimposed residues and close to zero to non-superimposed residues.

Next we examined the global quality estimates (Figure 2). The correlations between the global measures are higher than for the local measures. The correlations between the three superposition-based measures are excellent (*cc* > 0.95). In fact, the correlation between S-score and GDT_TS is astonishingly high (*cc* > 0.996) possibly explaining the success of the ProQ methods in CASP, where GDT_TS is used for model quality evaluation. The correlation between the two contact-based measures, CAD-score and lDDT, is also excellent (*cc* > 0.98), while the correlations between the contact-based and the superposition-based measures are lower (*cc* = 0.86 to 0.92).

**Figure 2:**
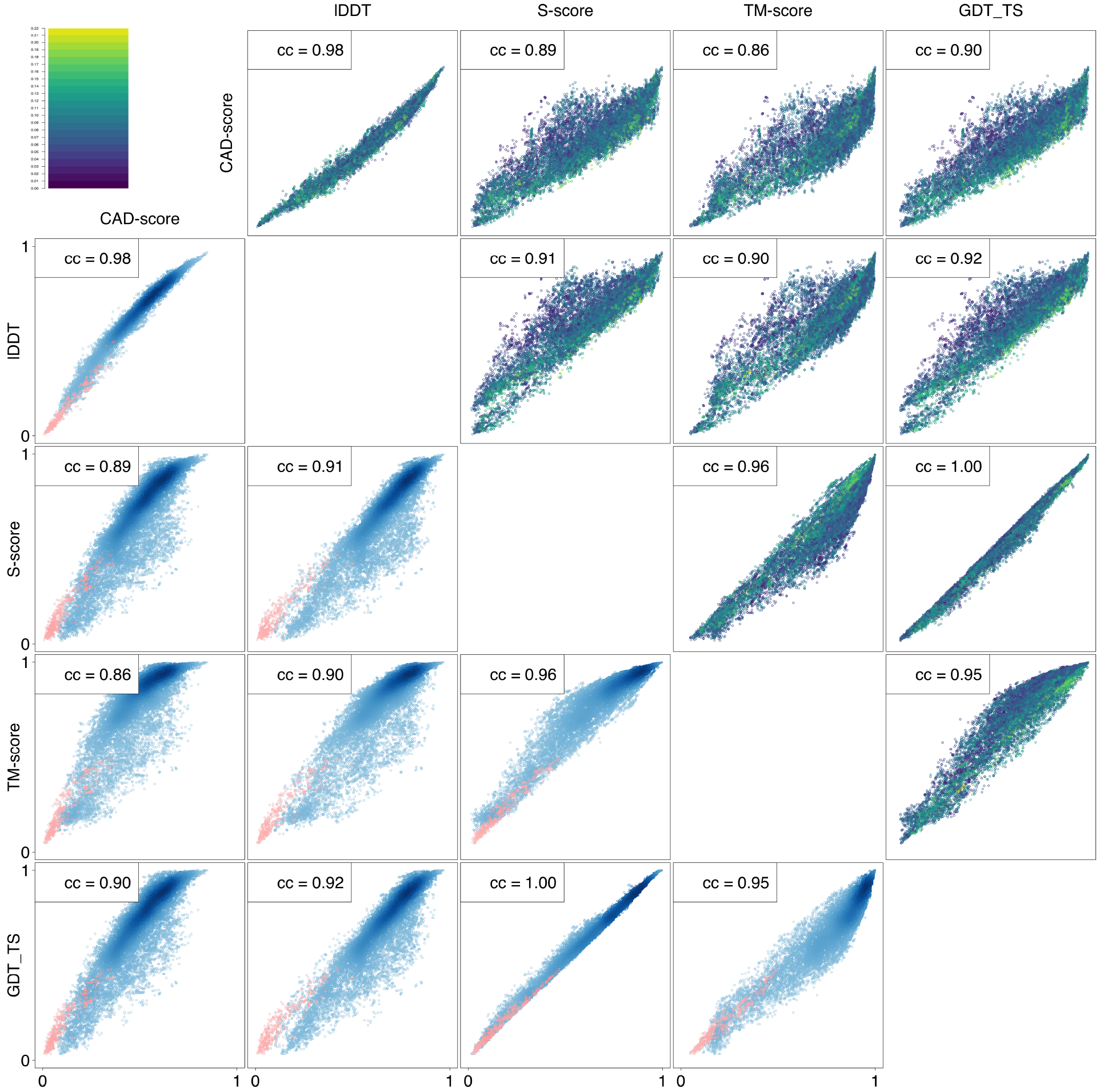
Scatter plot comparing different quality estimators on the *global* level using 19,985 models (points) submitted to CAMEO. The scatter plots on the upper right side are colored based on the contact order of the native structure, blue being low and yellow being high, see the color bar for actual values. The scatter plots on the lower left side are colored based on density of the points. The models that are shorter than 50% of the full length are colored in red. The correlation coefficient (cc) between the different quality estimators is given in the insets.

There are some differences in how the quality measures treat models that are not full length, i.e models that do not include all residues in the native structure. The models that are shorter than 50% of the full length are colored in red in Figure 2 and are often the outliers between different quality measures.

It can be noted that although the correlations are high between the different quality measures it is not higher than the correlation between predicted and estimated quality for the best quality predictors. In CASP12 the correlation of the consensus-based model quality predictors ^30, 31^ is higher to GDT_TS than the correlation between GDT_TS and CAD-score^10^.

### Training using different target functions

Next we set out to retrain ProQ3D using the four different quality measures as target functions. A deep learning network was trained on CASP9 and CASP10 models and evaluated on CAMEO (in main text) and CASP11 (see supplementary data). Below, the quality measures are referred to as ProQ3D-S, ProQ3D-TM, ProQ3D-CAD and ProQ3D-lDDT while the evaluation methods are referred to by their names (GDT_TS, S-score, TM-score, CAD-score and lDDT).

As expected, the best correlation is obtained when the same quality estimator is used both for training and testing (Figure 3). Clearly, on the local level it is easier to predict lDDT than any of the other measures. Independently, on the training method the correlation is always highest when using lDDT to evaluate the local quality.

**Figure 3:**
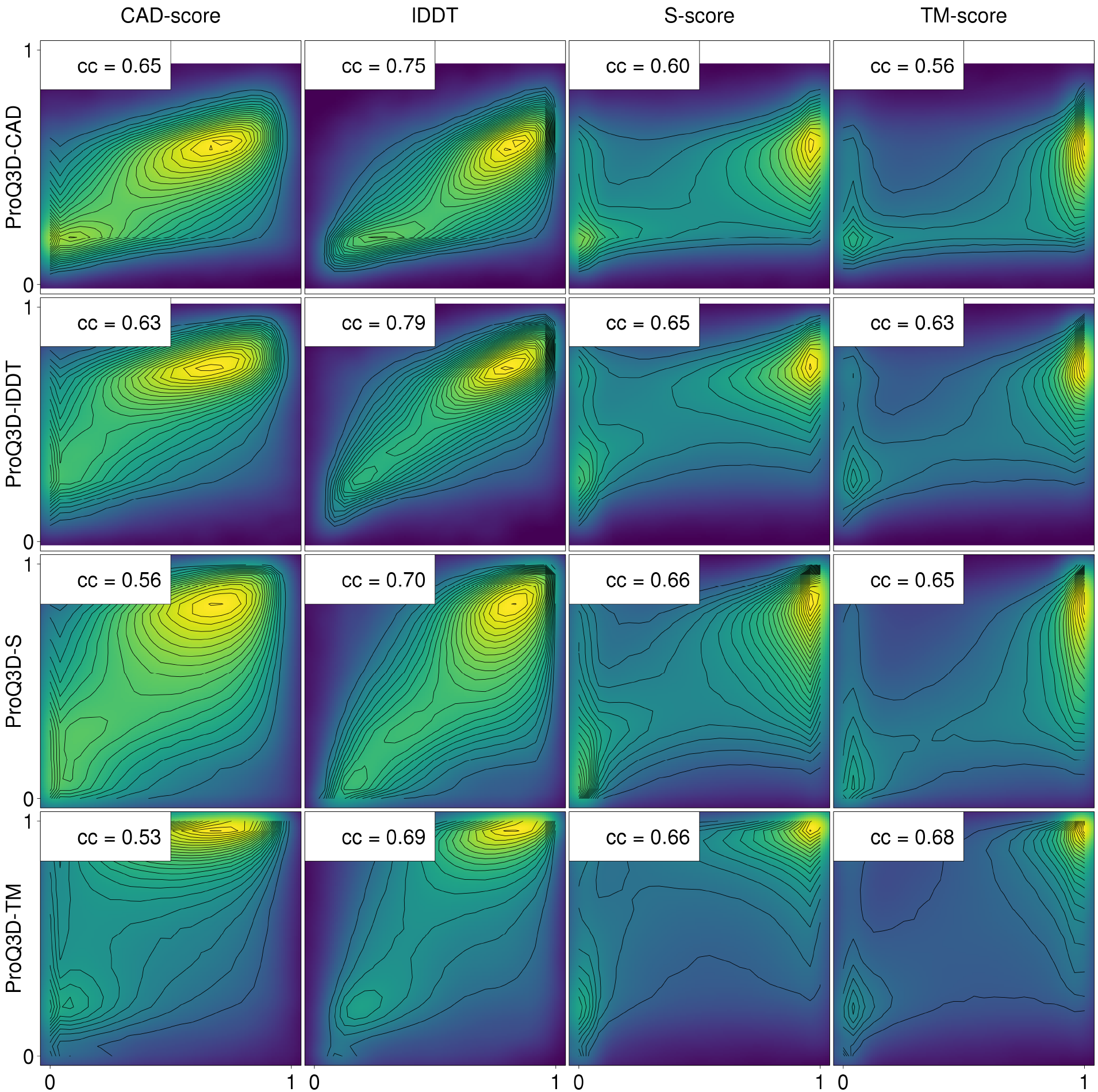
Scatter plot comparing real scores against the predicted scores on the *local* level using 4,939,249 residues (points) submitted to CAMEO. The intensity is plotted using the Viridis color scale, i.e. the most dense regions are yellow. Contour lines based on the point density are marked. The correlation coefficient (cc) between the different quality estimators is given in the insets.

After transferring the local predictions to global quality estimates it is still clear that the highest correlation is obtained when using the same training and evaluation function. However, the differences are smaller, with correlations in the range *cc* = 0.78 − 0.88, for all combinations of training and evaluation functions (Figure 4). Also, for global quality estimations both of the contact-based evaluations seems slightly easier to predict.

**Figure 4:**
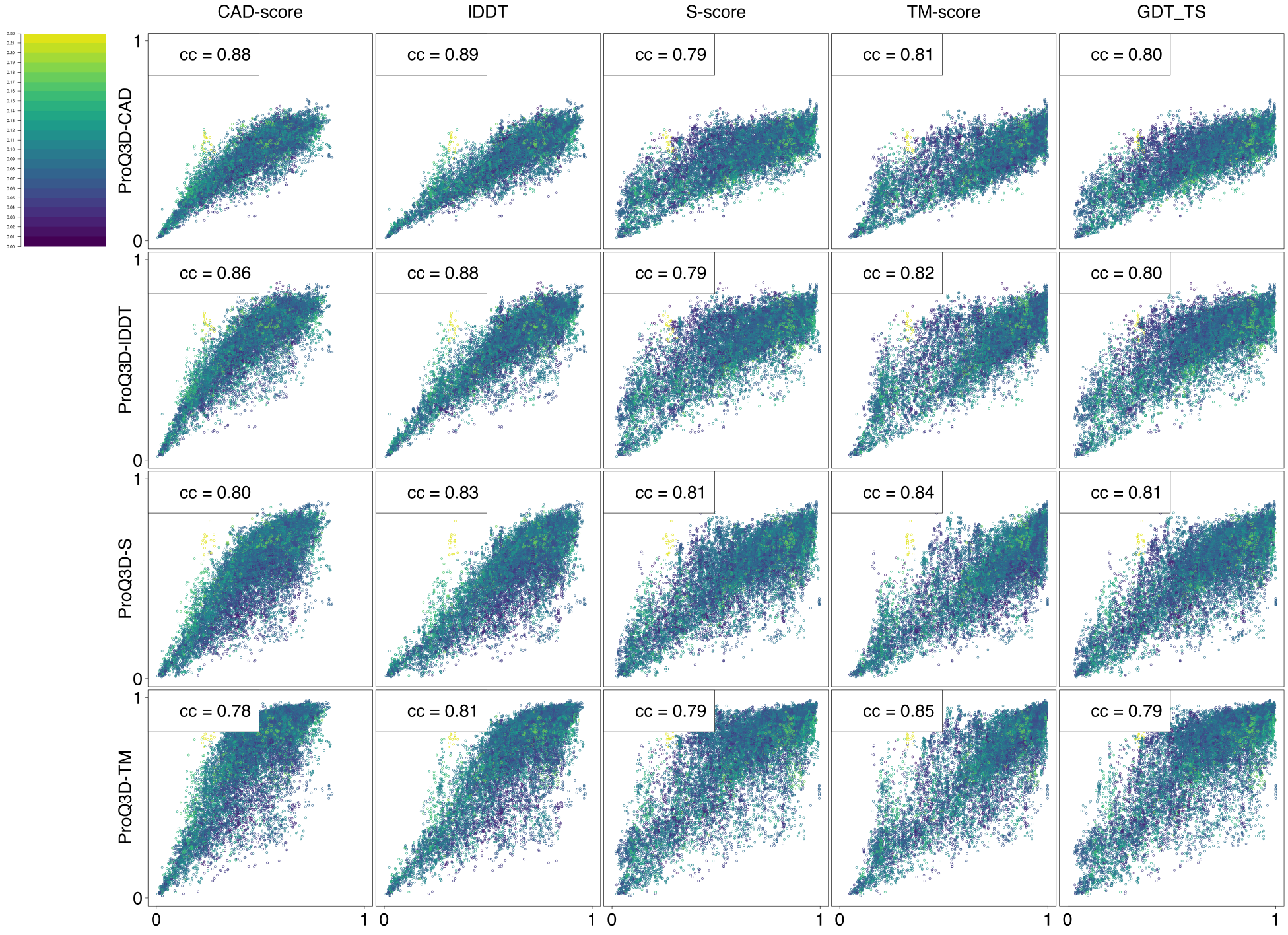
Scatter plot comparing real scores against the predicted scores on the *global* level using 19,985 models (points) submitted to CAMEO. The scatter plots are coloured based on the contact order of the native structure, blue being low and yellow being high, see the color bar for actual values. The correlation coefficient (cc) between the different quality estimators is given in the insets.

### Low contact order models differentiate the quality measures

Superposition-based quality estimators have problems with models that contain multiple domains, where the individual domains are correctly modeled but their relative arrangement is wrong. As mentioned above a superposition-based quality estimation methods superimpose one of the domains and assign low quality to all residues in the other domain. In contrast, the contact-based measures assign more appropriate domain level scores to the residues, if each domain is considered in isolation. It should be noted that this problem does not only relate to classical multi-domain proteins, but also to proteins with a fibrillar (or extended) shape, or actually any protein with a low contact-order. Therefore, we decided to study the targets with respect to their contact order in the native structure.

An example of a low contact order protein where CAD-score gives a much higher evaluation than S-score is CASP11 target T0821 shown Figure 5. This target has the lowest contact order in CASP11 data set (0.026) (see Figure S6) and almost all of the submitted models for this target have higher CAD-score than S-score, see red dots in Figure 5a. T0821 is a helical solenoid and most models are locally correct but the exact orientation of the N- and C-terminal parts is wrong. Evaluation of the model BAKER-ROSETTASERVER_TS4 highlights this. It has both units modelled more or less correctly, but their relative orientation is wrong. Therefore, S-score is only good for the part that is superposed (Figure 5b) while CAD-score gives relatively good scores to the entire protein (Figure 5c). As a result, the global CAD-score of the model is higher than its S-score.

**Figure 5:**
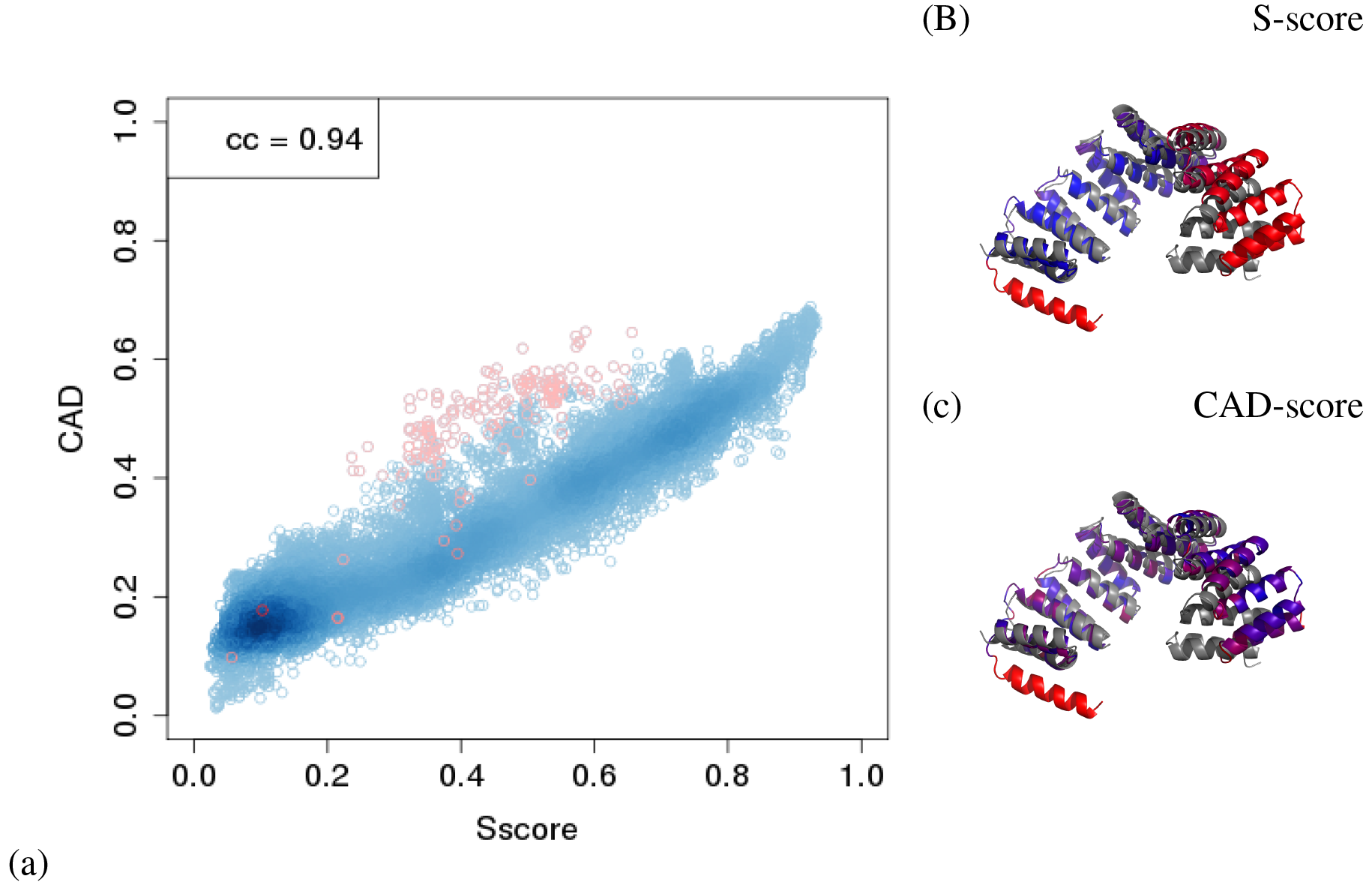
CAD-score and S-score for all models in CASP11. All models from target T0821 are marked in red. (b) BAKER-ROSETTASERVER_TS4 model for T0821 target coloured by S-score and superposed to the native structure. Good regions are coloured in blue, bad regions are coloured in red. The native structure is coloured in grey. The isolated helix that is coloured in red in the model is not present in the native structure (c) BAKER-ROSETTASERVER_TS4 model for T0821 target coloured by CAD-score. Good regions are coloured in blue, bad regions are coloured in red. The isolated helix that is coloured in red in the model is not present in the native structure

In Figure 4 and upper right side of Figure 2, the models are colored in blue to yellow according to their contact order, with low contact order targets being coloured in blue. When we compare the superposition-based scores (S-score and TM-score) with the two contact-based measures (CAD-score and lDDT) we can see that low contact order are more frequent to the upper left side of the diagonal. This means that the (blue) low contact order models achieve relatively higher qualities using the contact-based measures, as expected if only a part of the model has been superposed.

We can also notice that the S-score vs TM-score plot has more low contact order proteins on the lower right side of the diagonal. However, these differences rise mostly because of the different way how S-score and TM-score normalize the scores by protein length. TM-score gives higher score for longer proteins because of the way *d*0 value is calculated (see formula in Methods). Long proteins in turn tend to have lower contact order, despite the fact that we calculate a relative contact order which is normalized by protein length (see Supplementary Figure S6).

### Ranking models

It is not only important to have a good correlation for all protein models but also for each individual target. A good per target correlation enables the identification of the best model for that particular target. Here, the advantage of contact-based measures is clear, see Table I and Supporting Table S1. There are significantly higher self-correlations for lDDT and CAD-score (*cc* > 0.8 v s. *cc* < 0.6). Further, when training on a contact-based measures, the per target correlation is higher even when using the superposition-based evaluation methods on the CAMEO set, Table I, while this is not the case on the CASP set, Supporting Table S1.

**Table I:** Global predictions-average per target correlations on the CAMEO data set. Best prediction for each quality measure is marked in bold. Predictions that do not differ significantly (*p-value* < 0.001) from the best one in a column are also marked in bold. Wilcoxon signed-rank test was used to evaluate the significance. Please, note that according to Wilcoxon signed rank test a higher difference in the average value between two samples does not always mean that the difference is more significant.

| Predictor | CAD-score | IDDT | S-score | TM-score | GDT_TS |
| --- | --- | --- | --- | --- | --- |
| ProQ3D-CAD | **0.82** | **0.80** | 0.59 | 0.61 | 0.61 |
| ProQ3D-IDDT | 0.80 | **0.80** | **0.60** | **0.62** | **0.61** |
| ProQ3D-S | 0.61 | 0.64 | 0.53 | 0.55 | 0.54 |
| ProQ3D-TM | 0.66 | 0.69 | 0.55 | **0.59** | 0.57 |

Also the contact-based measures provide an improved detection of the best models as measured by the first ranked loss^9^, Table II. Training on CAD-score or lDDT selects higher quality models across all evaluation measures. Thus, using CAD-score or lDDT as target functions seem to provide a small but significant improvement in model selection. It can be noted that when training on a version of lDDT that only uses the *Cα*-atoms, lDDT-CA ^18^, the performance is very similar to the results obtained by lDDT. The results are slightly better at CASP11 and slightly worse using CAMEO, and for both datasets better than when using S-score or TM-score, see Supplementary Tables S1-4. Therefore, we do believe that the increased performance is mainly due to the difference between contact- and superposition-based quality evaluation functions and not due to the difference in what atoms are included in the analysis.

**Table II:**
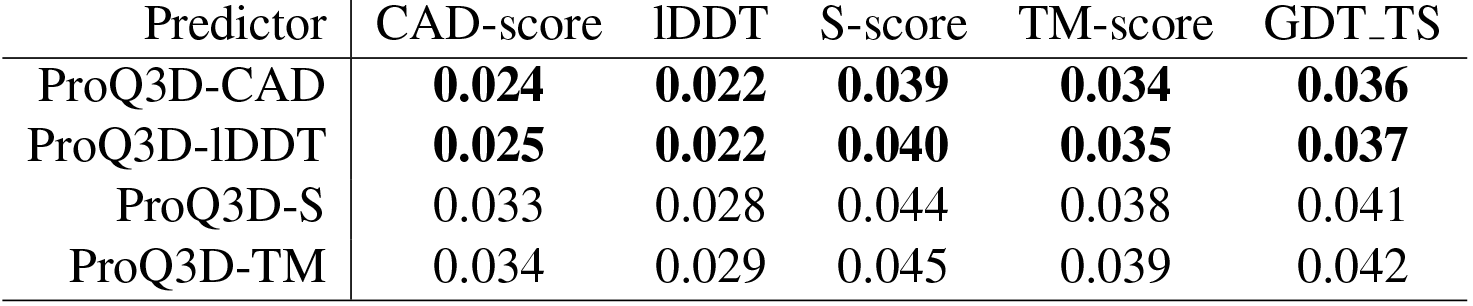
First ranked loss on the CAMEO data set. Best prediction for each quality measure is marked in bold. Predictions that do not differ significantly (*p-value* < 0.001) from the best one in a column are also marked in bold. Wilcoxon signed-rank test was used to evaluate the significance.

When studying GDT_TS for the models that are first ranked by different predictors it can be seen that for most targets models of very similar quality are selected (Figure 6). However, there appears to be a set of targets where the contact-based predictors consistently choose models that are significantly different from the ones selected by the superposition-based predictors. Four of these targets are analysed in more detail, see Table III.

**Table III:**
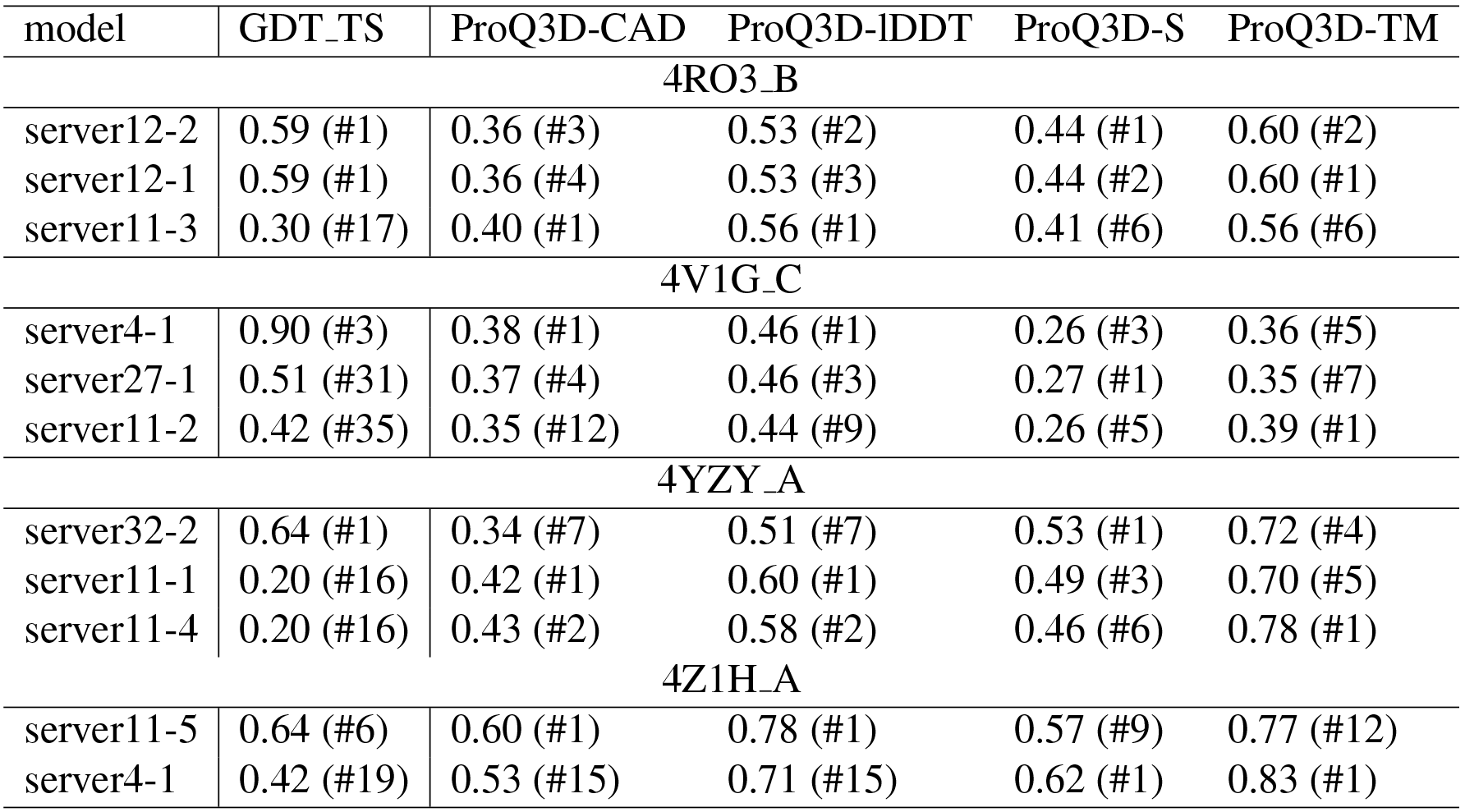
Four targets, where the top ranked models selected by the different ProQ3D methods have vastly different quality. The table shows the models that are top ranked by four different ProQ3D versions for four targets in CAMEO data set. GDT_TS score, as well as all four predicted scores are shown for each of these models. In parenthesis, the rank of the model is shown when sorted by the score of a particular predictor. In all and four examples ProQ3D-CAD and Proq3D-lDDT selects the same top ranked model.

**Figure 6:**
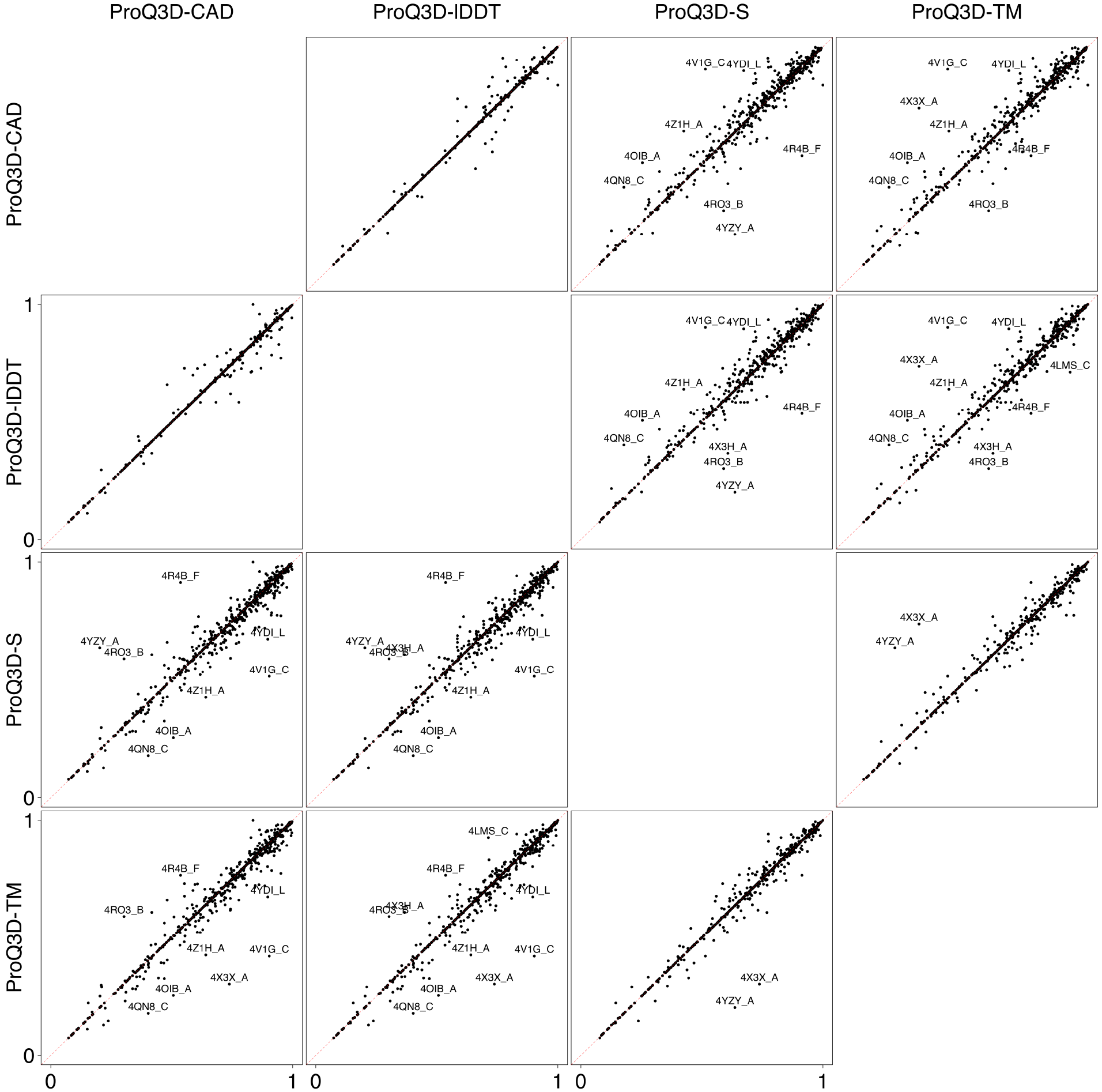
Scatter plot comparing GDT_TS of selected models by ProQ3D trained on different quality measures. CAMEO data set. Targets where GDT_TS difference of selected models is higher than 0.2 are labelled.

**Figure 7:**
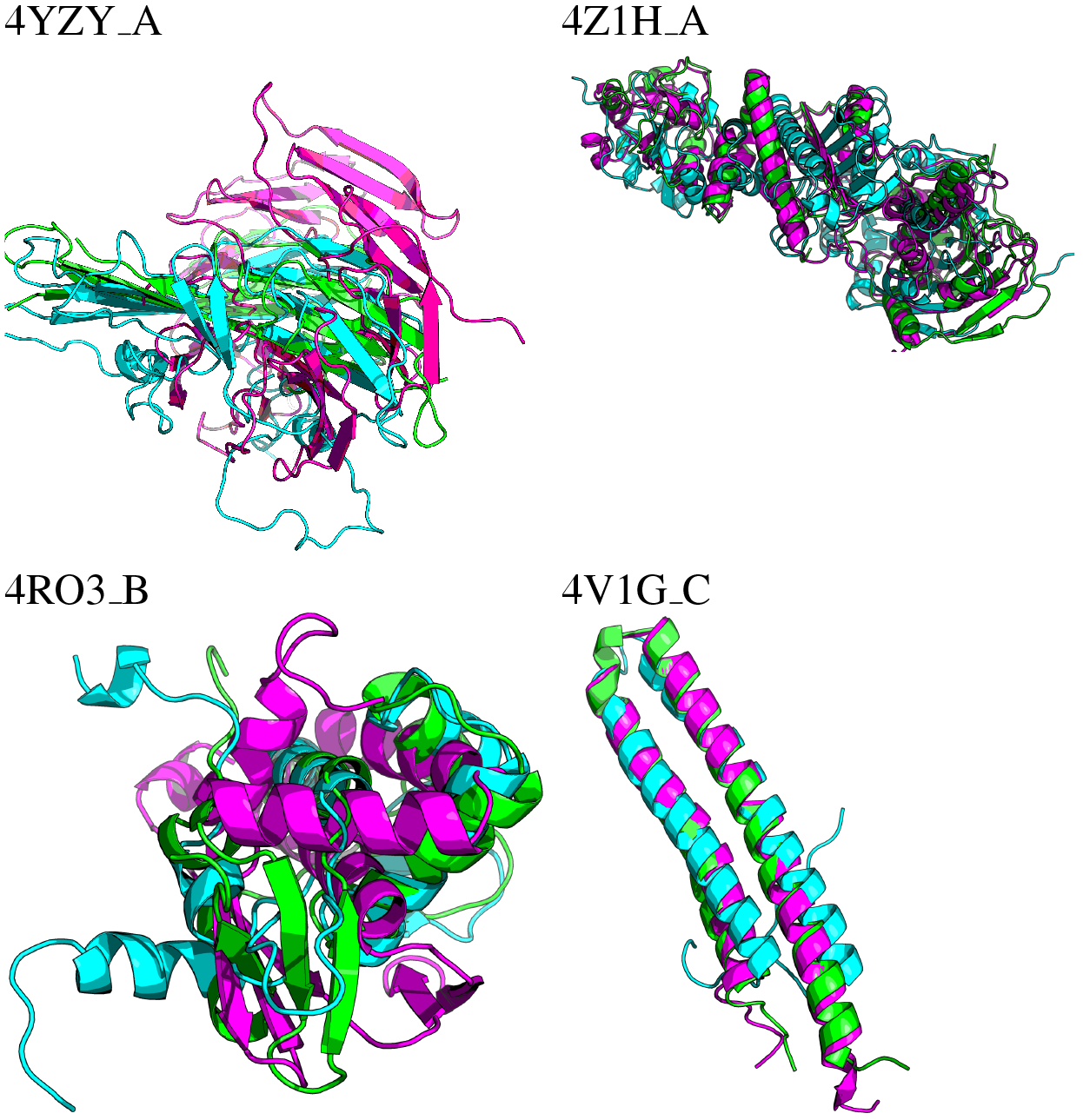
Top ranked models of target four tagets superposed to the native structure (green). Models selected by by ProQ3D-S are blue and by ProQ3D-CAD magenta.

For the target 4YZY_A, ProQ3D-S ranks the best model first, while the other predictors rank significantly worse models first. It can be seen in Figure 7 that the Proq3D-S model is very good while the the model selected by ProQ3D-CAD has a different fold. Also, for another target, 4RO3 B, both of the superposition-based predictors, ProQ3D-S and ProQ3D-TM, rank better models first than contact-based predictors. For the remaining two targets 4 V1G_C and 4 Z1H_A the contact-based predictors, ProQ3D-CAD and ProQ3D-lDDT, rank better models first.

In all of these four cases the difference in predicted scores between the good and the bad models are quite small. We did not find any obvious reason why one predictor ranked one model before the other one-indicating that when scores are similar and the models are different it is advisable to examine several models for a given target.

## Conclusions

Here, we show that when ProQ3D is retrained using different target functions, i.e. different descriptions of the model quality of a protein, model selection can be improved. Target functions can be divided into two groups: superposition- and contact-based methods. We find that when using contact-based measures as target functions in training, we can improve performance in model selection. We also notice that it is easier to predict the quality when using a contact-based description, in particular lDDT. However, the correlation with the “standard” CASP evaluation measure, GDT_TS, is best when the original superposition-based target function, S-score, is used for training ProQ3D.

## Funding

This work was supported by grants from the Swedish Research Council (VR-NT 2016-03798 to AE and 2012-5270 to BW) and Swedish e-Science Research Center (BW). The Swedish National Infrastructure provided computational resources for Computing (SNIC) at NSC.

